# Reducing gravity takes the bounce out of running

**DOI:** 10.1101/123745

**Authors:** Delyle T. Polet, Ryan T. Schroeder, John E. A. Bertram

**Affiliations:** Department of Biological Sciences, University of Calgary, Calgary T2N 1N4, Canada; Biomedical Engineering, University of Calgary, Calgary T2N 1N4, Canada; Cumming School of Medicine, University of Calgary, Calgary T2N 1N4, Canada

**Keywords:** bipedal running, reduced gravity, leg swing, energetics, optimization, biomechanics

## Abstract

In gravity below Earth normal, a person should be able to take higher leaps in running. We asked ten subjects to run on a treadmill in five levels of simulated reduced gravity and optically tracked center of mass kinematics. Subjects consistently *reduced* ballistic height compared to running in normal gravity. We explain this trend by considering the vertical takeoff velocity (defined as maximum vertical velocity). Energetically optimal gaits should balance energetic costs of ground-contact collisions (favouring lower takeoff velocity), and step frequency penalties such as leg swing work (favouring higher takeoff velocity, but less so in reduced gravity). Measured vertical takeoff velocity scaled with the square root of gravitational acceleration, following energetic optimality predictions and explaining why ballistic height decreases in lower gravity. The success of work-based costs in predicting this behaviour challenges the notion that gait adaptation in reduced gravity results from an unloading of the stance phase. Only the relationship between takeoff velocity and swing cost changes in reduced gravity; the energetic cost of the down-to-up transition for a given vertical takeoff velocity does not change with gravity. Because lower gravity allows an elongated swing phase for a given takeoff velocity, the motor control system can relax the vertical momentum change in the stance phase, so reducing ballistic height, without great energetic penalty to leg swing work. While it may seem counterintuitive, using less “bouncy” gaits in reduced gravity is a strategy to reduce energetic costs, to which humans seem extremely sensitive.

**Summary Statement:** During running, humans take higher leaps in normal gravity than in reduced gravity, in order to optimally balance the competing costs of stance and leg-swing work.

## Introduction

Under normal circumstances, why do humans and animals select particular steady gaits from the myriad possibilities available? One theory is that the chosen gaits minimize metabolic energy expenditure (Alexander and Jayes, 1983; Ruina et al., 2005). To test this theory, one can subject organisms to abnormal circumstances. If the gait changes to a new energetic optimum, it can be inferred that energetics also govern gait choice under normal conditions (Bertram and Ruina, 2001; Long and Srinivasan, 2013; Selinger et al., 2015).

One “normal” gait is the bipedal run, and one abnormal circumstance is that of reduced gravity. Movie 1 demonstrates the profound effect reducing gravity has on running kinematics. A representative subject runs at 2 m s^−1^ in both Earth-normal and simulated lunar gravity (about one-sixth of Earth-normal). The change in kinematics is apparent; the gait in normal gravity involves pronounced center-of-mass undulations compared to the near-flat trajectory of the low-gravity gait. While center-of-mass vertical excursions during stance are known to decrease in reduced gravity (Donelan and Kram, 2000), we observed that the height achieved in the flight phase also decreases. This gait modification seems paradoxical: in reduced gravity, people are free to run with much higher leaps. Instead, they seem to flatten their gait. Why should this be?

A simple explanation posits that the behaviour is energetically beneficial. To explore the energetic consequences of choosing to run with lower leaps in reduced gravity, we first considered the impulsive model of running, following Rashevsky (1948) and Bekker (1962), which treats a human runner as a point mass body bouncing off rigid vertical limbs (Fig. 1). Stance is treated as an inelastic, impulsive collision with the ground. In reality, stance occurs in finite time, and elastic mechanisms exist. However, the inelastic approximation is remarkably productive in explaining gait choice (Ruina et al., 2005). When we use the term “energetic cost of collisions”, we are generally referring to non-recoverable energy loss during stance resulting from some interaction of the center of mass with the ground (Bertram and Hasaneini, 2013). Such losses may arise from damping, active negative work or discontinuous velocity profiles. In any case, modelling these interactions as an inelastic collision provides a simple estimation of the net cost.

**Figure 1:**
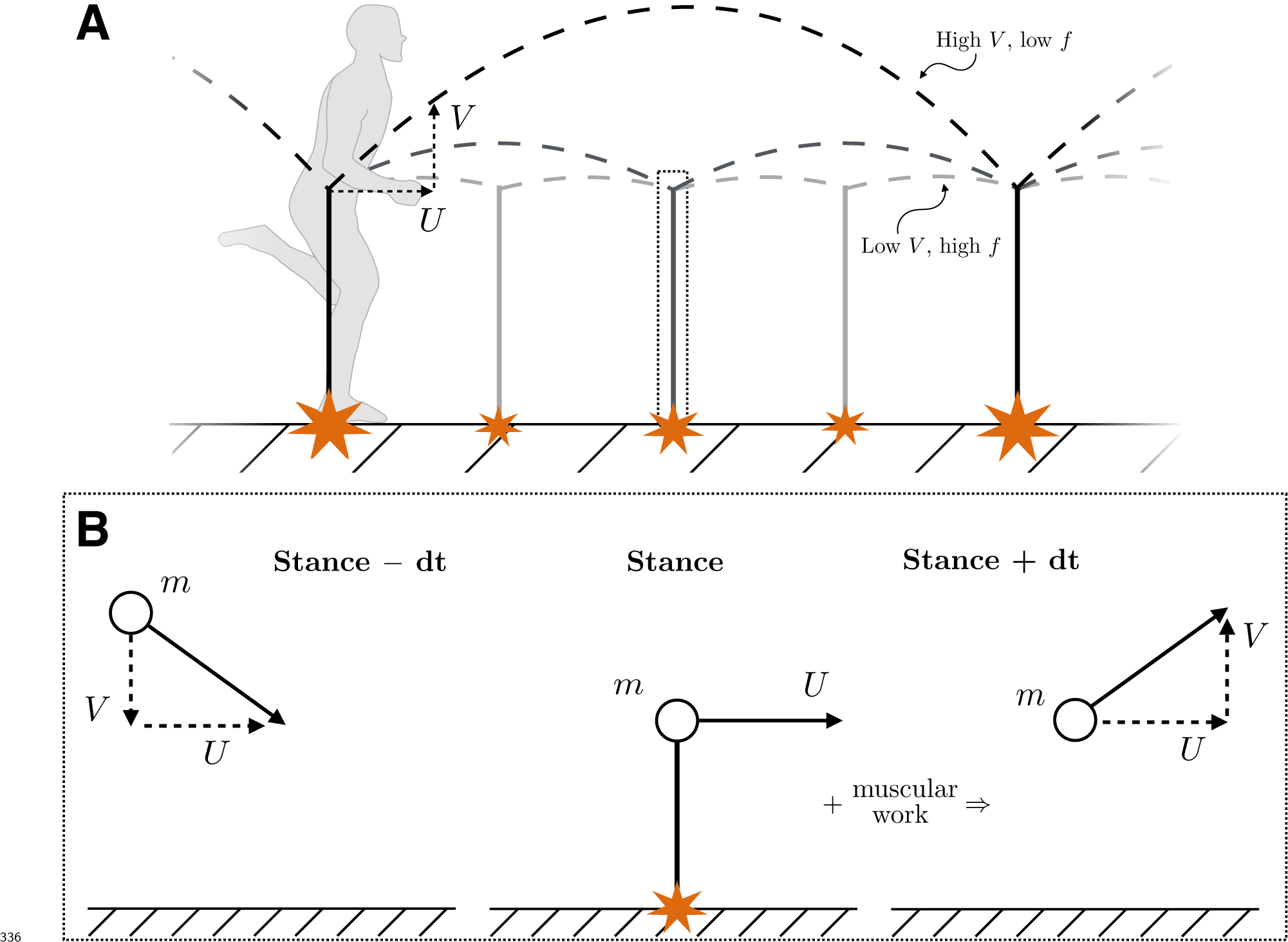
Schematics explaining the energetic model. (A) In the impulsive model of running, a point mass bounces off vertical, massless legs during an infinitesimal stance phase. As the horizontal velocity *U* is conserved, the vertical takeoff velocity *V* dictates the step frequency and stride length. Smaller takeoff velocities (light grey) result in more frequent steps that incur an energetic penalty, while larger takeoff velocities (dark grey) reduce the frequency penalty but increase losses during stance. The small box represents a short time around stance that is expanded in panel B. (B) We assume that the center-of-mass speed at landing is equal to the takeoff speed. The vertical velocity *V* and its associated kinetic energy are lost during an impulsive foot-ground collision of infinitessimally short duration. The lost energy must be resupplied through muscular work. Horizontal acceleration is assumed small and is neglected in the model.

During this collision, all vertical velocity is lost while horizontal velocity is conserved (Fig. 1b). The total kinetic energy lost per step is therefore *E_col_* = *mV*^2^/2, where *m* is the runner’s mass and *V* is their vertical takeoff velocity^1^. Lost energy must be recovered through muscular work to maintain a periodic gait, and so an energetically-optimal gait will minimize these losses. If center-of-mass kinetic energy loss were the only source of energetic cost, then the optimal solution would always be to minimize vertical takeoff velocity. However, such a scenario would require an infinite stepping frequency as *V* approaches zero (Alexander, 1992; Ruina et al., 2005), as step frequency (ignoring stance time and air resistance) is *f* = *g*/(2*V*), where *g* is gravitational acceleration.

Let us suppose there is an energetic penalty that scales with step frequency, as *E*_freq_ ∝ *f^k^* ∝ *g^k^* /*V^k^*, where *k* > 0. Such a penalty may arise from work-based costs associated with swinging the leg, which are frequency-dependent (*k* = 2; Alexander, 1992; Doke et al., 2005), or from short muscle burst durations recruiting less efficient, fast-twitch muscle fibres (*k* ≈ 3; Kram and Taylor, 1990; Kuo, 2001). Notably, this penalty increases with gravity, since the non-contact duration will be shorter for any given takeoff velocity in higher gravity. The penalty also has minimal cost when *V* is maximal; smaller takeoff velocities require more frequent steps, which is costly. Therefore, the two sources of cost act in opposite directions: collisional loss promotes lower takeoff velocities, while frequency-based cost promotes higher takeoff velocities.

If these two effects are additive, then it follows that the total cost per step is

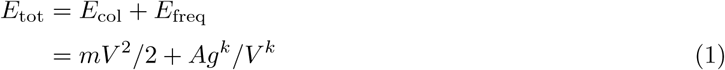

where *A* is an unknown proportionality constant relating frequency to energetic cost. As the function is continuous and smooth for *V* > 0, a minimum can only occur either at the boundaries of the domain, or when 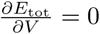. Solving the latter equation for *V* yields

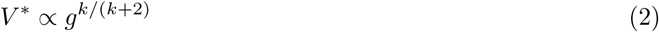

as the unique critical value. Here the asterisk denotes a predicted (optimal) value. Since *E_tot_* approaches infinity as *V* approaches 0 and infinity (Eqn 1), the critical value must be the global minimum in the domain *v*> 0. As *k* > 0, it follows from Eqn 2 that the energetically-optimal solution is to reduce the vertical takeoff velocity as gravity decreases.

The observation^2^ of He et al. (1991) that 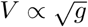 implies *k* = 2, a finding consistent with frequency costs arising from the work of swinging the limb (Alexander, 1992; Doke et al., 2005). However, their empirical assessment of the relationship used a small sample size, with only four subjects. We tested the prediction of Eqn 2 by measuring the maximum vertical velocity over each running stride, as a proxy for takeoff velocity, in ten subjects using a harness that simulates reduced gravity. We also measured the maximum vertical displacement in the ballistic phase to verify whether the counter-intuitive observation of lowered ballistic center-of-mass height in hypogravity, as exemplified in Movie 1, is a consistent feature of reduced gravity running.

## Materials and Methods

We asked ten healthy subjects to run on a treadmill for two minutes at 2 m s^−1^ in five different gravity levels (0.15, 0.25, 0.35, 0.50 and 1.00 G, where G is 9.8 m s^−2^). A belt speed of 2 m s^−1^ was chosen as a comfortable, intermediate jogging pace that could be accomplished at all gravity levels. Reduced gravities were simulated using a harness-pulley system similar to that used by Donelan and Kram (2000), but differing in the use of a spring-pendulum system to generate near-constant force for a large range of motion. Hasaneini et al. (2017 preprint) provide more details of the apparatus. The University of Calgary Research Ethics Board approved the study protocol and informed consent was obtained from all subjects. Leg length for each subject was measured during standing from the base of the shoe to the greater trochanter on one leg.

Due to the unusual experience of running in reduced gravity, subjects were allowed to acclimate at their leisure before indicating they were ready to begin each two-minute measurement trial. In each case, the subject was asked to simply run in any way that felt comfortable. Data from 30 to 90 s from trial start were analyzed, providing a buffer between acclimating to experimental conditions at trial start and initiating slowdown at trial end.

### Implementation and measurement of reduced gravity

Gravity levels were chosen to span a broad range. Of particular interest were low gravities, at which the model predicts unusual body trajectories. Thus, low levels of gravity were sampled more thoroughly than others. The order in which gravity levels were tested was randomized for each subject, so as to minimize sequence conditioning effects.

For each gravity condition, the simulated gravity system was adjusted in order to modulate the force pulling upward on the subject. In this particular harness, variations in spring force caused by support spring stretch during cyclic loading over the stride were virtually eliminated using an intervening lever (see Figs 3 and 4 in Hasaneini et al., 2017, preprint). The lever moment arm was adjusted in order to set the upward force applied to the harness, and was calibrated with a known set of weights prior to all data collection. A linear interpolation of the calibration was used to determine the moment arm necessary to achieve the desired upward force, given subject weight and targeted effective gravity. Using this system, the standard deviation of the upward force during a trial (averaged across all trials) was 3% of the subject’s Earth-normal body weight.

Achieving exact target gravity levels was not possible since the lever’s moment arm is limited by discrete force increments (approximately 15 N). Thus, each subject received a slight variation of the targeted gravity conditions, depending on their weight. A real-time data acquisition system allowed us to measure tension forces at the gravity harness and calculate the effective gravity level at the beginning of each new condition. The force-sensing system consisted of an analog strain gauge (Micro-Measurements CEA-06-125UW-350), mounted to a C-shaped steel hook connecting the tensioned cable and harness. The strain gauge signal was passed to a strain conditioning amplifier (National Instruments SCXI-1000 amp with SCXI-1520 8-channel universal strain gauge module connected with SCXI-1314 terminal block), digitized (NI-USB-6251 mass termination) and acquired in a custom virtual instrument in LabView. The tension transducer was calibrated with a known set of weights once before and once after each data collection trial to correct for modest drift error in the signal. The calibration used was the mean of the pre- and post-experiment calibrations.

### Center of mass kinematic measurements

A marker was placed at the lumbar region of the subject’s back, approximating the position of the center of mass. Each trial was filmed at 120 Hz using a Casio EX-ZR700 digital camera. The marker position was digitized in DLTdv5 (Hedrick, 2008). Position data were differentiated using a central differencing scheme to generate velocity profiles, which were further processed with a 4^th^-order low-pass Butterworth filter at 7 Hz cutoff. The vertical takeoff velocity was defined as the maximum vertical velocity during each gait cycle (*V* in Fig. 1). This definition corresponds to the moment at the end of stance where the net vertical force on the body is null, in accordance with a definition of takeoff proposed by Cavagna (2006).

Vertical takeoff velocities were identified as local maxima in the vertical velocity profile. Filtering and differentiation errors occasionally resulted in some erroneous maxima being identified. To rectify this, first any maxima within ten time steps of data boundaries were rejected. Second, the stride period was measured as time between adjacent maxima. If any stride period was 25% lower than the median stride period or less, the maxima corresponding to that stride period were compared and the largest maximum kept, with the other being rejected. This process was repeated until no outliers remained.

Position data used to determine ballistic height were processed with a 4^th^-order low-pass Butterworth filter at 9 Hz cutoff. Ballistic height was defined as the vertical displacement from takeoff to the maximum height within each stride. No outlier rejection was used to eliminate vertical position data peaks, since the filtering was slight and no differentiation was required. If a takeoff could not be identified prior to the point of maximum height within half the median stride time, the associated measurement of ballistic height was rejected; this strategy prevented peaks from being associated with takeoff from a different stride.

### Statistical methods

Takeoff velocities and ballistic heights were averaged across all gait cycles in each trial for each subject. To test whether ballistic height varied with gravity, a linear model between ballistic height and gravitational acceleration was fitted to the data using least squares regression, and the validity of the fit was assessed using an *F*-test. A linear model was also tested for log(*V*) against log(*g*) using the same methods. Since the proportionality coefficient between *V*\* and 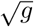 is unknown *a priori*, we derived its value from a least squares best fit of measured vertical takeoff velocity against the square root of gravitational acceleration, setting the intercept to zero. Given a minimal correlation coefficient of 0.5 and sample size of 50, a *post-hoc* power analysis yields statistical power of 0.96, with type I error margin of 0.05. Data were analyzed using custom scripts written in MATLAB (v. 2016b).

## Results

### Response of Ballistic Height and Takeoff Velocity to Gravity

Data from all trials are shown in Fig. 2. Ballistic height increases with gravity (Fig. 2A, linear vs constant model, *p* = 4 × 10^−4^, *R*^2^ = 0.24, *N* = 50), validating the counter-intuitive result exemplified in Movie 1 as a consistent feature of running in hypogravity.

**Figure 2:**
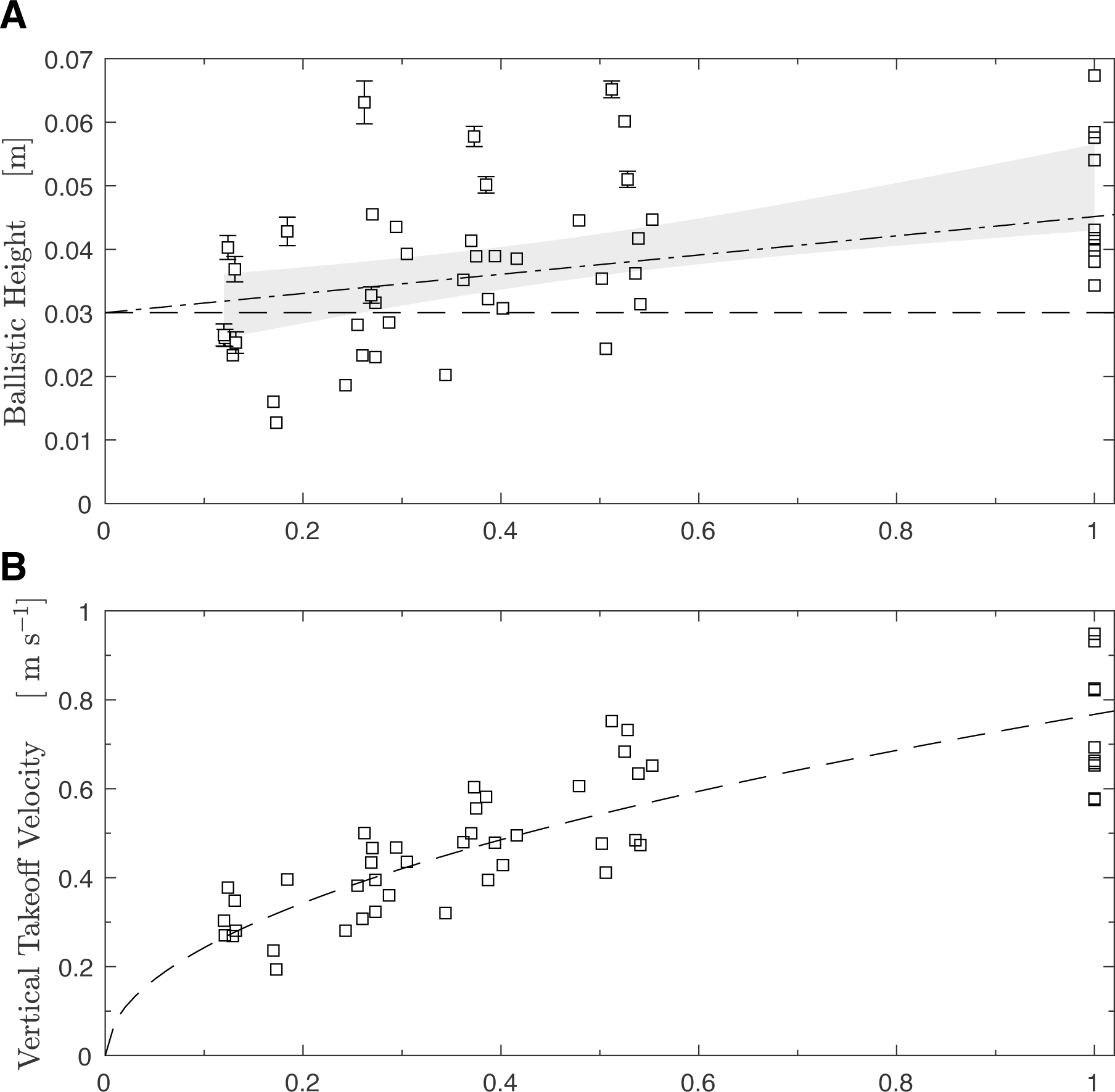
Human subjects lower both ballistic height and takeoff velocity during running in reduced gravity. (A) Mean ballistic height (data points) increases with gravity (*p* of linear vs constant model under two-tailed *F*-test: 4 × 10^−4^, *N* = 50). The dashed line is the prediction for ballistic height from the impulsive model, which deviates from observation at high *g*. The dash-dot line adds a correction factor for finite stance time from the spring mass model (Eqn 3). This second prediction lies within the 95% CI of the least-squares linear fit (grey area). Both predictions use takeoff velocities from the best fit in panel B. (B) Measured vertical takeoff velocities increase proportionally with the square root of gravitational acceleration, following work-based energetic optimality. The least squares fit of the impulsive model with *k* = 2 is shown as a dashed line. The fit has an *R*^2^ value of 0.73 (*N* = 50). Each data point is a mean value measured in one subject (ten subjects total) across multiple steps (*n* ≥ 50) during a one-minute period at a given gravity level. For both panels, if error bars (twice the s.e.m.) are smaller than the markers, then they are not shown. Data used for creating these graphics are given in Table S1.

Takeoff velocity also increases with gravitational acceleration (Fig. 2B), and a least-squares fit of Eqn 2 using *k* = 2 follows empirical measurements well (*R*^2^ = 0.73, *N* = 50). Other values of *k* were also tested (Fig 3). If the impulsive model is accurate, then the best-fit slope of a scatter plot of log(*V*) against log(*g*) should correspond to *k*/(*k* + 2) (Eqn 2), that is, slopes of 0.33, 0.50 or 0.60 for *k* = 1, 2 and 3 respectively. Only the slope predicted by *k* = 2 falls within the 95% confidence interval of the least squares slope (0.47 ± 0.09; Fig. 3).

**Figure 3:**
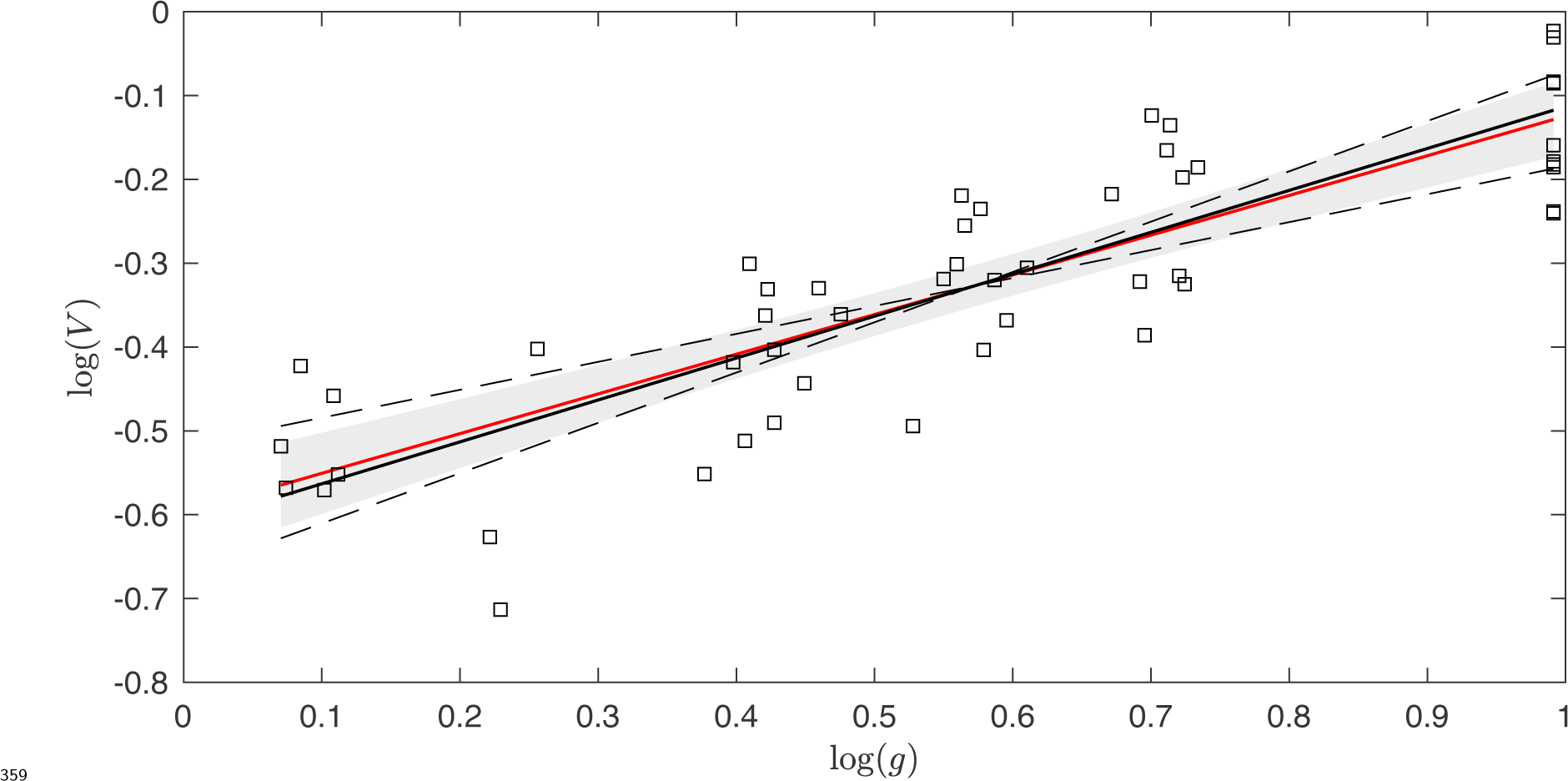
A log-log plot of vertical takeoff velocity against gravitational acceleration shows that the impulsive model yields the best fit when *E*_freq_ ∝ *f*^2^. The least squares linear fit is shown in red as a solid line, with 95% confidence interval as a grey area. The linear fit exhibits *R*^2^ = 0.70 and a slope of 0.47±0.09 (best estimate ± 95% CI, *N* = 50), which is not significantly different from the predicted slope of 0. 5 for *k* = 2 (black solid line), where *k* is the exponent relating frequency to cost (*E*_freq_ ∝ *f^k^*). Both *k* =1 and *k* = 3 (shallow and steep dashed lines, respectively) yield predicted slopes (0.33 and 0.60, respectively) that lie outside the 95% CI, indicating that a work-based swing cost at *k* = 2 is a superior fit to the data, while a simple linear frequency cost (*k* = 1) and an approximate force/time cost (*k* = 3, see Kuo 2001) do not represent these data well. Data points are from ten subjects running at five gravity conditions each, and each point is the mean of at least 64 takeoffs measured during each trial.

A best fit at *k* =2 implies a frequency-based cost arising primarily from the work of leg swing. However, since only the center of mass is offloaded by the harness, the natural frequency of limb swing remains unchanged for all target gravity levels (Donelan and Kram, 2000). Since metabolic energy of swing is minimal at natural frequency (Doke et al., 2005), it is necessary to adjust the predictions from the impulsive model (Appendix A). An adjusted model exhibits a fit with *R*^2^ =0.745 (*N* = 50, Fig. A1), only marginally better than the simple model with *k* = 2 (*R*^2^ = 0.73, Fig. 2B). The predictions do not change greatly, because time spent in the air *is* affected by gravity, and more air time requires less work to swing the legs, regardless of natural frequency^3^.

**Figure A1:**
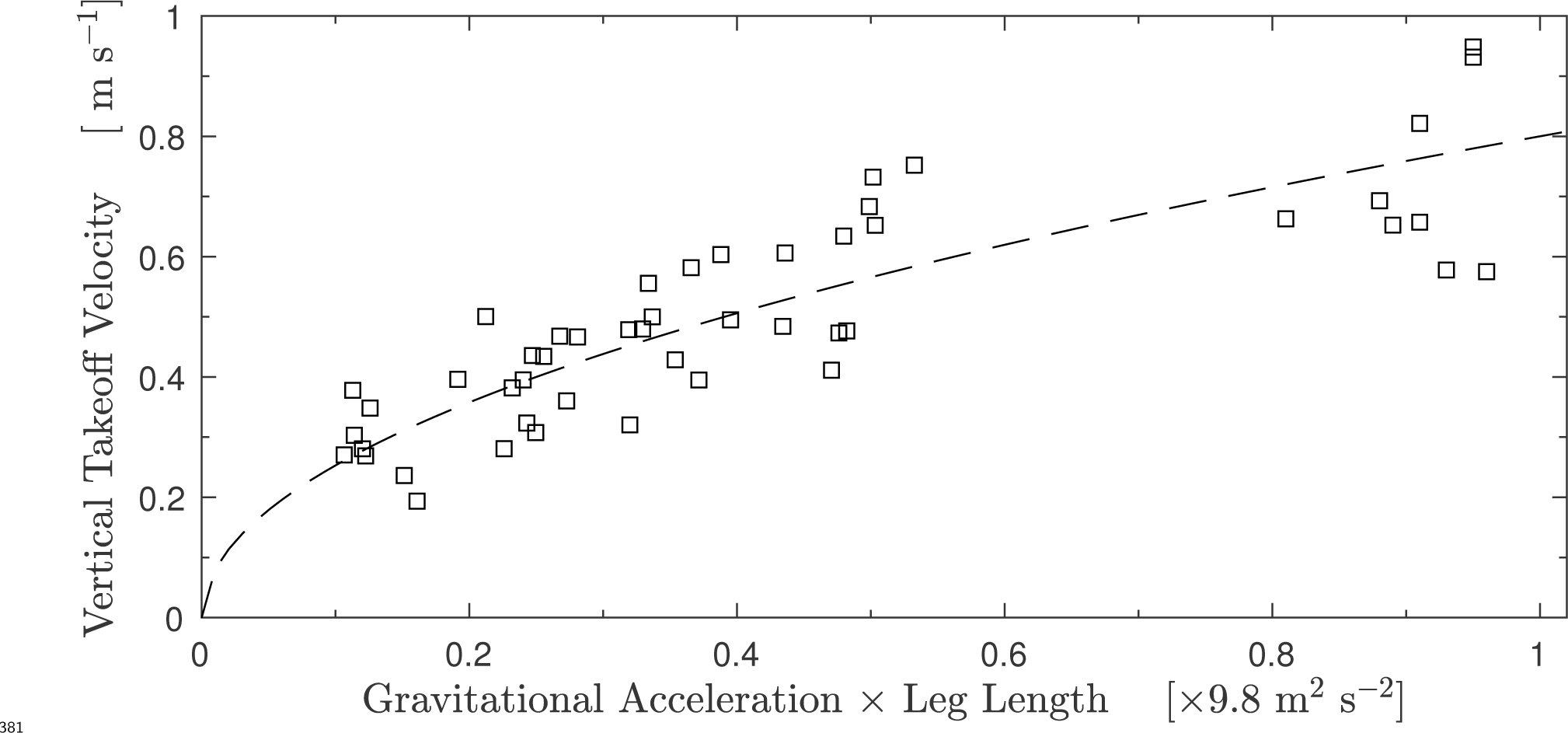
Vertical takeoff velocity scales with the square root of gravitational acceleration times leg length during running. The least squares fit for the model given by Eqn A4 is shown as a dashed line. The fit exhibits *R*^2^= 0.745, using all fifty data points. Error bars (twice standard error of the mean takeoff velocity measured during a trial, *n* ≥ 64) are smaller than the marker size.

### Predicting Ballistic Height Trends

The impulsive model with *k* = 2 predicts that the ballistic height should remain constant (dash line in Fig. 2A). This constant value agrees with empirical data at low *g*, but exhibits increasing error towards normal *g*.

We defined “takeoff” as occurring when the net force on the body was null and velocity was maximal; however, this does not equate to the moment when the stance foot leaves the ground. After the point of maximal velocity, upward ground reaction forces decay to zero. During this time, the net downward acceleration on the body is less than gravitational acceleration. Thus, the body travels higher than would be expected if maximal velocity corresponded exactly to the point where the body entered a true ballistic phase, as in the model (Fig. 1).

We can account for the missing impulse with the spring-mass model. This model describes the kinematics and dynamics of running well (McMahon and Cheng, 1990; He et al., 1991; Blickhan and Full, 1993), and provides a way to estimate stance time from takeoff velocity (though it lacks the ability to *predict* takeoff velocity; McMahon and Cheng, 1990). Notably, correcting the prediction 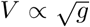 with spring-mass model estimates of finite stance yields the following relationship for ballistic height (Appendix B):

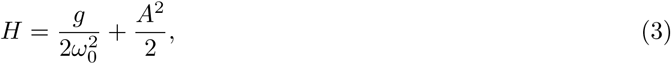

where *ω*_0_ is the natural angular frequency of vertical oscillation, and *A* is a constant in the relationship 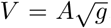 Note that Eqn 3 is linear in *g*, and approaches the predictions from the impulsive model alone as *g* → 0. Taking *ω*_0_ ≈ 18 rad s^−1^ from He et al. (1991), and A from the best-fit in Fig. 2B, Eqn 3 gives the dot-dash line shown in Fig. 2A. The predicted relationship (Eqn 3) has a slope of 0.015 m G^−1^ and an intercept of 0.03 m, and is within the 95% confidence interval of the best-fit slope (0.021 ± 0.01 m G^−1^) and intercept (0.029 ± 0.006 m), indicating that finite stance accounts for the discrepancy within error, though it underpredicts the true slope somewhat.

## Discussion

Human runners lower the height achieved in the ballistic phase as gravity decreases. This adaptation requires pronounced modification of the takeoff velocity, since maintaining the latter parameter in all conditions would result in substantially increased ballistic height in reduced gravity. Why human runners would modify their gait so greatly was initially unclear.

A simple work-based model of energetic cost explains the trends well. The fit in Fig. 2B exhibits an *R*^2^ value of 0.73, indicating that a simple energetic model can explain over two thirds of the variation in maximum vertical velocity resulting from changes in gravity. Human runners seem to be sensitive to these energetic costs and adjust their takeoff velocity accordingly. However, the model has its limitations, and an accounting of finite stance (which was initially neglected in the model) was necessary to explain the trend of increasing ballistic height with gravity. Despite the updated model matching the general trend of the data, the slope in Eqn 3 is reduced compared to the empirically-derived slope.

The use of the external lumbar point as a center of mass approximation may explain some of the remaining difference between Eqn 3 and observation. At lower gravity, the body maintained a relatively erect, rigid posture (as exemplified by Movie 1), and so the lumbar marker likely follows the center of mass closely. However, at higher gravity, the legs move through larger excursions and the torso exhibits slight rotation, making the lumbar estimate less accurate. At normal gravity, Slawinski et al. (2004) showed that the lumbar point overestimates vertical oscillations of the flight phase (by less than 1 cm)– though their trials were at a high belt speed (5 m s^−1^). If the same results hold in our case, we would expect that the measured ballistic height in Fig. 2A should be slightly lower at higher levels of gravity, reducing the actual slope and possibly improving the agreement to Eqn 3. Future work could use a multisegment model to improve center of mass and ballistic height measurements, but such a technique is unlikely to reverse the trend of increasing ballistic height with gravitational acceleration.

The present results indicate that the cost of step frequency is a key factor in locomotion. Although the exact value of the optimal takeoff velocity depends on both frequency-based penalties and collisional costs, the former penalties change with gravity while the latter do not (Fig. 4). The collisional cost landscape is independent of gravity because the final vertical landing velocity is alone responsible for the lost energy. Regardless of gravitational acceleration, vertical landing speed must equal vertical takeoff speed in the model; so a particular takeoff velocity will have a particular, unchanging collisional cost.

**Figure 4:**
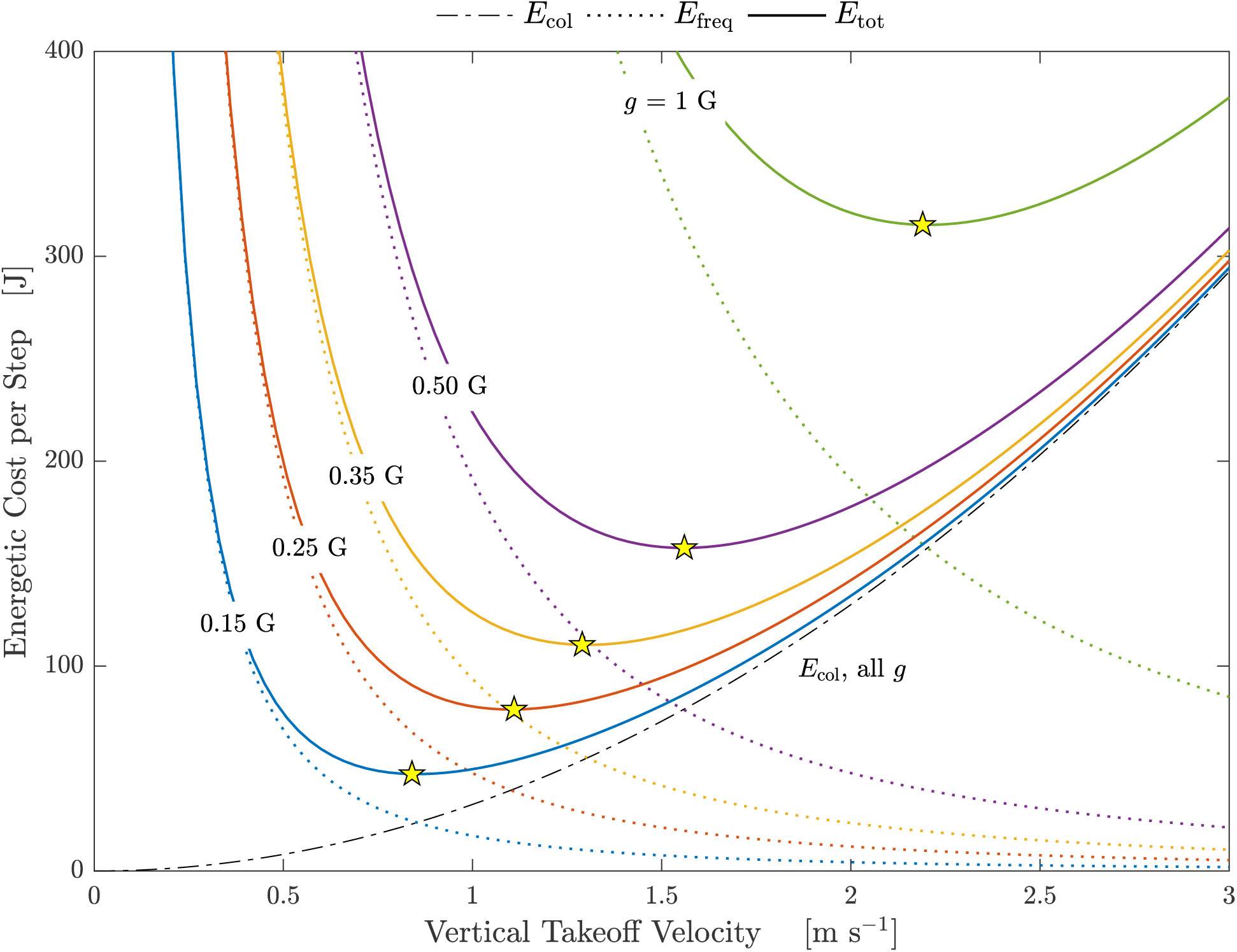
The energetic costs according to the model are plotted as a function of vertical takeoff velocity (*V*) for the five levels of gravity tested. The hypothetical subject has a mass of 65 kg and a frequency-based proportionality constant (A in *E*_freq_ = *Af*^2^) derived from the best fit in Fig. 2B. Labels of gravity levels (g) are placed over the colours they represent. The collisional cost curve (*E*_col_ = *mV*^2^/2, black dot-dash line) does not change with gravity, while the frequency-based energetic cost curve (*E*_freq_, dotted lines) is sensitive to gravity, leading to an effect on total energy per step (*E*_tot_, solid lines). In lower gravity, a runner can stay in the air longer for a given takeoff velocity, so the associated frequency-based cost goes down. However, the cost of collisions at that same velocity is unchanged, since it depends only on the velocity itself. The relaxation of frequency-based cost allows the runner settle on a lower, optimal takeoff velocity (yellow stars) with both a lower frequency-based and collisional cost, compared to higher gravity.

However, taking off at a particular vertical velocity results in greater flight time at lower levels of gravity; thus, the frequency-based cost curves are decreased as gravity decreases (Fig. 4). Frequency-based costs, particularly limb-swing work, appear to be an important determinant of the effective movement strategies available to the motor control system. Their apparent influence warrants further investigation into the extent of their contribution to metabolic expenditure.

While the present study corroborates others in finding that a work-based cost (*k* = 2) predicts locomotion well (Alexander, 1980, 1992; Hasaneini et al., 2013), other authors have favoured a higher-order “force/time” cost (Kuo, 2001; Doke et al., 2005; Doke and Kuo, 2007). Interestingly, a higher-order model in frequency cost (*k* = 3) did not fit the present data; however, our simple model with *k* = 3 only approximates the force/time cost in the swing phase, and does not account for a rate cost during stance. Further research must be done to distinguish the predictive value of work-based cost to its alternatives; however, for the present results, a work-based model is sufficient, at least for takeoff velocity.

The present results challenge the notion that metabolic cost of running is determined largely by the cost of generating force during stance (Kram and Taylor, 1990; Arellano and Kram, 2014), purportedly supported by the observation that metabolic cost is proportional to gravity (Farley and McMahon, 1992). According to the best-fit model presented here, the net cost (Eqn 1) at optimal takeoff velocity (Eqn 2) is expected to increase in proportion to gravitational acceleration (that is, *E*_tot_(*V*\*) ∝ *g*), as Farley and McMahon observed (1992). The cost of vertical acceleration of the center of mass can decrease as gravity is reduced *only* because the relationship between takeoff velocity and swing cost changes; this allows the subject to settle on a lower stance cost, whose relationship to takeoff velocity does *not* change as a function of gravity (Fig. 4). These trends can be explained simply from muscular work, and do not rely on any independent force-magnitude cost.

The model presented in this article is admittedly simple and makes unrealistic assumptions, including impulsive stance, no horizontal muscular force, non-distributed mass, and a simple relationship between step frequency and energetic cost. Further, horizontal accelerations will incur a larger portion of energetic losses as horizontal speed increases (willems et al., 1995), and the tradeoff between swing and stance costs may change. The present model would not be able to anticipate any such trend, as it has no dependence on horizontal speed. Future investigations could evaluate work-based costs using more advanced optimal control models (Srinivasan and Ruina, 2006; Hasaneini et al., 2013), eliminating some of these assumptions and allowing for an investigation into horizontal speed dependence. Despite its simplicity, the impulsive model with work-based swing cost is able to correctly predict the observed trends in takeoff velocity with gravity, and demonstrates that understanding the energetic cost of both swing and stance is critical to evaluating why the central nervous system selects specific running motions in different circumstances.

Although many running conditions are quite familiar, running in reduced gravity is outside our general experience. Surprisingly, releasing an individual from the downward force of gravity does not result in higher leaps between foot contacts. Rather, humans use less bouncy gaits with slow takeoff velocities in reduced gravity, taking advantage of a reduced collisional cost while balancing a stride-frequency penalty.

## Acknowledgements

The authors would like to thank Art Kuo, Jim Usherwood, David Lee and Allison Smith for comments on earlier drafts, as well as two anonymous reviewers who provided constructive insights that greatly improved the manuscript.

## Author Contributions

All authors assisted in designing the experiment, collecting data and writing the manuscript; D.T.P. conceived the energetics-based model and performed data analysis. All authors gave final approval for submission.

## Data Availability

The dataset supporting this article has been uploaded as part of the supplementary material (Table S1).

## List of Symbols

*θ*: leg angle (radians)
*ω*_0_: vertical natural angular frequency in the spring-mass model (radians s^−1^)
*A*: proportionality constant in the relationship *E*_freq_ = *Af^k^* (J s^k^)
*B*: proportionality constant in the relationship 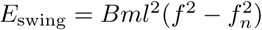
*E*_col_: energetic cost of collisions (J)
*E*_freq_: energetic cost related to step-frequency (J)
*E*_swing_: energetic cost of leg swing work (J)
*E*_tot_: total energetic cost (*E*_col_ + *E*_freq_ or *E*_col_ + *E*_swing_, in J)
*f*: step frequency (Hz)
*f_n_*: natural pendular frequency (Hz)
*g*: gravitational acceleration (m s^−2^)
*G*: Earth-normal gravitational acceleration (9.8 m s^−2^)
*Gr*: Groucho number (≡ *νw*_0_/*g*)
*H*: ballistic height (m)
*I*: leg moment of inertia about the hip (kg m^2^)
*k*: exponent in proportionality *E*_freq_ ∝ *f^k^*
*l*: leg length (m)
*m*: total subject mass (kg)
*r*: length change from leg rest length (m)
*t*: time after toe-down (s)
*t*\*: time at which maximum vertical speed is achieved (s)
*t_m_*: time at which maximum vertical velocity is achieved (s)
*t_s_*: stance period (s)
*U*: average horizontal speed (m s^−1^)
*ν*: vertical velocity at toe-off (m s^−1^)
*V*: vertical velocity at takeoff (maximum vertical velocity,in m s^−1^)
*V*\*: optimal and predicted vertical takeoff velocity (m s^−1^)

## Appendices

### A Cost of swing work in partial reduced gravity

The experimental apparatus (Hasaneini et al., 2017, preprint) unloads a subject’s center of mass, but does not act directly on their limbs. Consequently, while their center of mass might experience reduced weight, the limbs swing under the influence of normal gravity. It is prudent to check how this affects the predictions of the impulsive model.

The work required to swing a limb is (Doke et al., 2005)

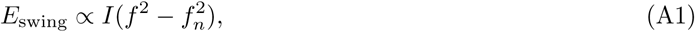

where *I* is the moment of inertia of the limb about the hip, *f* is the frequency of oscillation and *f_n_* is the natural frequency (equal to 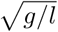 for a simple pendulum, where *l* is leg length). Here we are assuming that the limb changes configuration little during the swing phase, and so *I* is approximately constant. Note that Eqn A1 is only valid when *f* > *f_n_* (Doke et al., 2005), since if sufficient time is available the limb can swing passively. The swing frequency is slightly greater than the stride frequency^4^, which in the present study ranged from trial-mean values of 0.69 to 1.47 Hz over all subjects and conditions (Table S1). Doke et al. (2005) found the natural frequency of swinging legs to be 0.64 ± 0.02 Hz (mean ± s.d.) for a subject group with mean leg length of 0.88 ± 0.07 m (mean ± s.d., *N* = 12). Our subject group exhibited larger mean leg length (0.92 ± 0.06 m, mean ± s.d., *N* = 10), so would very likely have smaller natural frequencies. Therefore, the assumption that *f* > *f_n_* very likely holds in this case.

The leg moment of inertia about the hip scales approximately as *I* ∝ *ml*^2^, where *m* is body mass and *l* is the leg length (Winter, 2009). Assuming *f* = *g*/(2*V*), and invoking *E*_tot_ = *E*_col_ + *E*_swing_, we have

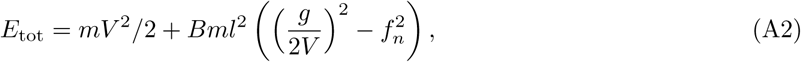

where *B* is some proportionality constant. To achieve the energetically optimal takeoff velocity, we take the derivative of Eqn A2 with respect to *V*, yielding

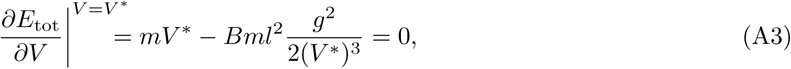

where we note that any dependence on *f_n_* has disappeared. However, there is a new dependence on *l*. Solving Eqn A3 for *V*\*, we find

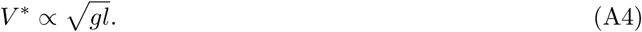

Empirical *V* is plotted against *gl* in Fig. A1 with the least square fit of Eqn A4. The fit exhibits *R*^2^ =0.745,only a slight improvement compared to the simple impulsive model (*R*^2^ = 0.73). Eqn A4 depends on *l*, but if the variation in *l* is small, then Eqn A4 is indistinguishable from the simple swing-cost model (Eqn 2 with *k* = 2). Indeed, the leg lengths of our subject group varied only by a factor of 1.3 (range 0.81 to 1.04 m), while the highest experimental *g* was six times the smallest value. Since the variation in leg length was comparatively small, it has little impact on the results.

### B Ballistic height corrections from the spring-mass model

We seek to predict the vertical center-of-mass displacement achieved between takeoff (maximum vertical velocity) and the maximum height during the flight phase. We know the maximum height from ballistics to be *v*^2^/(2*g*), where *v* is the vertical velocity at toe-off. However, we do not know the displacement between takeoff and toe off, nor do we know how to relate the velocity at takeoff to the velocity at toe-off. Both of these unknowns could be calculated using the ground reaction force during stance, but this was not measured empirically.

Instead, we can rely on the spring-mass model, which gives a decent approximation of the ground reaction forces assuming the velocity at toe-off and natural angular frequency (*ω*_0_) are given (McMahon and Cheng, 1990). In our case, the toe-off velocity is unknown, but the spring-mass model allows us to relate it to the maximum vertical velocity, which can in turn be predicted by the impulsive model. *ω*_0_ is defined as 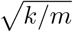 where *k* is the “spring” stiffness and *m* is mass. *k* is not actually the tendon stiffness, but is the virtual stiffness generated by the motor control system during stance (Farley and Ferris, 1998; Donelan and Kram, 2000); that is, the muscle and tendon forces combine to generate ground reaction forces as if there were one linear spring acting on the center of mass. The complicated interplay between muscles, tendons and energetics makes the angular frequency hard to predict.

Fortunately, the vertical spring stiffness is held more-or-less constant through changes in gravity (He et al., 1991), so we can use the empirically derived value^5^ of *ω*_0_ ~ 18 rad s^−1^. It remains simply to find the displacement between takeoff and toe-off, and the vertical toe-off velocity, in terms of the vertical takeoff velocity and gravity.

We follow McMahon and Cheng (1990) in assuming a point-mass body of mass *m* and massless legs. We assume that the ground reaction force is well-approximated by the compression of a spring with angular frequency *ω*_0_. For simplicity, we use a hopping model, which assumes that a person exhibits a small excursion angle (i.e. *θ* ~ 0). The leg length minus resting length is *r*, and so the dynamics of the system are

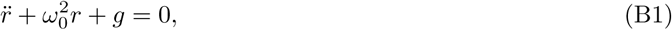

where *g* is gravitational acceleration. Setting the vertical landing velocity to *ṙ*(0) = −*ν*, and the initial position as *r*(0) = 0, the solution to the ordinary differential equation is (McMahon and Cheng, 1990)

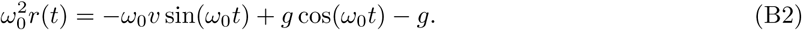

The instantaneous velocity is thus

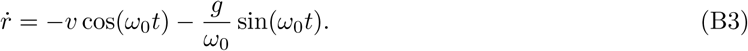

Eqns B1-B3 are valid for 0 ≤ *t* ≤ *t_s_*, where *t_s_* = (2π — 2arctan(Gr))/*ω*_0_ is the stance period, and we have introduced the non-dimensional Groucho number Gr ≡ *νω*_0_/*g* (McMahon and Cheng, 1990). For *t_s_* < *t* < *t_s_* + 2*ν*/*g*, the body is in a ballistic phase.

We can now determine the timing and magnitude of the peak vertical velocity. Let *t*\* correspond to any time at which a maximum speed is acheived. Since Eqn B3 is continuous and periodic, local maxima and minima in velocity must satisfy 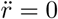. Therefore, from Eqn B1,

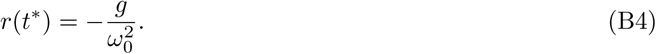

Combining Eqn B4 with B2 and solving for 0 ≤ *t*\* ≤ *t_s_* yields

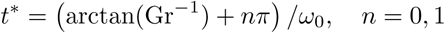

corresponding to the points of maximal speed during stance. The second point (*n* = 1), corresponds to the time at which maximal velocity is achieved,

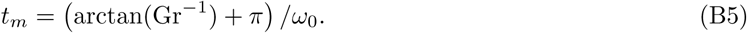

To determine the peak velocity *V*, we insert Eqn B5 into B3. Using the relations 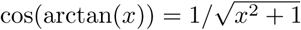 and 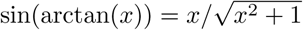, we find

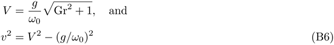

In the main manuscript, we define the ballistic height (*H*) as the vertical displacement from the time of maximal vertical velocity to the maximum height achieved during a stride, that is,

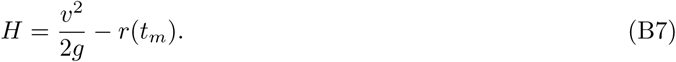

We need only insert Eqns B4 and B6 into Eqn B7 to find

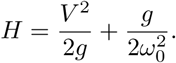

Note that the first term is identical to the prediction of the impulsive model (*i.e. V* = *ν*), while the second term gives a correction from the spring mass model, due to finite stance time. Since we have established that 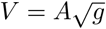 the prediction for *H* in terms of *g* alone is

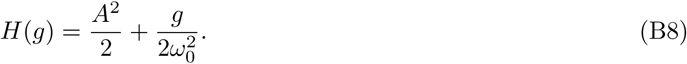

## Footnotes

1 In this paper, we are taking the vertical takeoff velocity as the maximum vertical velocity during the gait cycle, following Cavagna (2006). However, this is not always equal to the vertical velocity at take-off, and this distinction complicates the analysis. These complications are addressed in the discussion.

2 In reality, He et al. (1991) measured vertical speed at initial foot contact, but for the impulsive model in its simplest form, this is indistinguishable from takeoff velocity.

3 As long as stride frequency is greater than natural frequency, which is very likely the case for the present study (Appendix A)

4 Swing period is two flight phases and one stance phase, or one stance phase shorter than stride period.

5 This value was calculated by taking the average value of vertical stiffness data in Fig. 7 of He et al. (1991), dividing by average mass of subjects in the same study and taking the square root. 18 rad s^−1^ falls within all the error bars of Fig. 7, so seems representative of the natural angular frequency at all levels of gravity.

